# The human Toll-like receptor 2 (TLR2) response during pathogenic *Leptospira* infection

**DOI:** 10.1101/2023.11.16.567338

**Authors:** CN Kappagoda, RMISK Senevirathne, D Jayasundara, YPJN Warnasekara, LASM Srimantha, LAPNF De Silva, SB Agampodi

## Abstract

**Background:** Human innate immune responses are triggered through the interaction of human pattern recognition receptors and pathogen-associated molecular patterns. The role of toll-like receptor2 (TLR2) in mice innate immune response to leptospirosis is well established, while human studies are limited. The present study aimed to determine the TLR2 response among confirmed cases of leptospirosis.

**Methodology/Principle findings:** The study has two components. Clinically suspected patients of leptospirosis were confirmed using a previously validated qPCR assay. Total RNA was extracted from patients’ RNA-stabilized whole blood samples. Human TLR2 gene expression (RT-qPCR) analysis was carried out using an exon-exon spanning primer pair, using CFX Maestro™ software. The first set of patient samples was used to calculate the Relative Normalized Expression (ΔΔCq value) of the TLR2 gene in comparison to a healthy control sample and normalized by the reference gene GAPDH (Glyceraldehyde-3-phosphate dehydrogenase). Secondly, recruited patient samples were subjected to TLR2 gene expression analysis and compared to healthy controls and normalized by the reference genes Beta-2-microglobulin(B2M), Hypoxanthine phosphoribosyltransferase 1 (HPRT 1).

In the initial cohort of 64 confirmed leptospirosis cases, 18 were selected for human TLR2 gene expression analysis based on criteria of leptospiremia and RNA yield. Within this group, one individual exhibited a down-regulation of TLR2 gene (Expression/ΔΔCq=0.01352), whereas the remaining subjects presented no significant change in gene expression. In a subsequent cohort of 23 confirmed cases, 13 were chosen for similar analysis. Among these, three patients demonstrated down-regulation of TLR2 gene expression, with Expression/ΔΔCq values of 0.86574, 0.47200, and 0.28579, respectively. No TLR2 gene expression was noted in the other patients within this second group.

**Conclusions:** Our investigation into the acute phase of leptospirosis using human clinical samples has revealed a downregulation of TLR2 gene expression. This observation contrasts to the upregulation commonly reported in the majority of in-vitro and in-vivo studies of *Leptospira* infection. These preliminary findings prompt a need for further research to explore the mechanisms underlying TLR2’s role in the pathogenesis of leptospirosis, which may differ in clinical settings compared to laboratory models.

**Author Summary:** The human immune system employs pattern recognition receptors like toll-like receptor 2 (TLR2) to detect and combat infections such as leptospirosis. While TLR2’s role is well-documented in mice, its function in the human response to leptospirosis remains unclear. Our study evaluated TLR2 activity in patients with confirmed leptospirosis. We conducted a genetic analysis of blood samples from these patients, comparing TLR2 gene activity against healthy individuals, with standard reference genes for accuracy. Contrary to expectations and existing laboratory data, we observed a decrease in TLR2 activity in some patients. This suggests that human TLR2 responses in actual infections may diverge from established laboratory models. These findings indicate a need for further study to understand the human immune response to leptospirosis, which may significantly differ from that observed in controlled experimental settings.

## Introduction

The etiological agent responsible for Leptospirosis, namely *Leptospira*, was initially isolated a century ago through independent and concurrent efforts by Stimson and Inada. [1,2]. *Leptospira* spp. are thin, spiral, highly motile spirochetes that can be visualized by darkfield microscopy. Human leptospirosis is contracted by direct or indirect contact with leptospires in the urine of reservoir hosts, primarily via environmental interactions, owing to its zoonotic nature. Leptospirosis manifests as a wide range of clinical presentations from mild to severe disease [3]. The pathophysiology of human leptospirosis remains characterized by numerous areas of uncertainty. This is primarily attributed to the extensive diversity of *Leptospira* pathogens and the wide spectrum of clinical presentations associated with the disease. Therefore, further exploration is warranted to elucidate the pathophysiological mechanisms underlying leptospirosis.

The human body’s innate immune responses to pathogens are instigated through the recognition of Pathogen Associated Molecular Patterns (PAMPs) by a diverse array of Pattern Recognition Receptors (PRRs). PAMPs are expressed on the outer cell membrane of pathogens and are crucial for their survival and virulence. In the case of pathogenic *Leptospira* spp, lipopolysaccharides (LPS) and proteins have been identified as the principal PAMPs [4–7]. While there are various types of PRRs present in innate immune cells during *Leptospira* spp. infection, Toll-like receptors (TLRs) have garnered significant attention in the global scientific literature and are among the most extensively studied PRRs [8].

Of the TLRs studied, lipoprotein recognition of the pathogen has been suggested as a main role of TLR2. This notion was substantiated by demonstrating the higher susceptibility of TLR2-deficient mice in response to the gram-positive bacterium *Streptococcus pneumoniae*, as compared to wild-type mice [9]. The major outer membrane protein of *L.interrogans,* LipL32 [10], demonstrated as the main lipoprotein expressed during the mammalian leptospirosis infection, using hamster kidney tissues and clinical leptospirosis serum [11]. Wertz and colleagues demonstrated that both mice and human monocytic cells’ TLR2 receptors can initiate an innate immune response by recognizing *L.interrogans* LipL32 and LPS in the presence of the CD14 factor [12]. Additionally, further investigations utilized an atomic force microscope to evaluate the direct interaction between TLR2 and the cell surface receptors of the bacterium, providing conclusive evidence of TLR2’s interaction with the LipL32 receptors expressed on the surface of *Leptospira* [13]. The interaction of pathogenic *Leptospira* spp. with human PRRs triggers a response involving the secretion of immune mediators, cytokines, antimicrobial peptides, and the induction of the recruitment of other leukocytes aimed at eliminating the bacteria [14,15]. Conversely, studies have suggested that the impaired recognition of PAMPs by PRRs can enable the pathogen to evade the host’s innate immune responses, ultimately contributing to the development of more severe disease outcomes. Post-translational modifications of lipoproteins have altered the immune cell response to evade the host immunity [16]. Single nucleotide polymorphism(SNP) of TLR2(Arg753Gln) changed the functions of TLR2, which leads to disease development and the host lethality [17]. Although it was previously reported that LipL32 is expressed on the leptospiral cell surface, a recent study has suggested that LipL32 is localized beneath the outer cell membrane 18] This altered localization of LipL32 might result in its evasion from recognition by PRRs, contributing to the development of the disease.

While in-vitro and animal models have provided insights into the role of TLR2 in *Leptospira* infection, studies examining innate immune responses and the contribution of TLR2 in human leptospirosis are scarce. Hence, our study aimed to evaluate the messenger RNA (mRNA) level response of the human TLR2 gene among confirmed cases of leptospirosis by employing a novel exon-exon spanning primer pair for a reliable and efficient RT-qPCR assay.

## Methods

### Ethical statement and the study setting

All patients and healthy controls provided written informed consent for the study. The ethical approval for the research proposal was granted by the Ethics Review Committee, Faculty of Medicine and Allied Sciences, Rajarata University of Sri Lanka (ERC/2015/18). The present study was a part of a multi-center study on leptospirosis[19]. This study was confined to clinically suspected cases of leptospirosis admitted to the Teaching Hospital, Anuradhapura, located in the dry zone of Sri Lanka within the North Central Province.

### Enrollment of patients and healthy individuals

Clinically suspected leptospirosis patients for the present study were selected from a large ongoing project, in which all acute undifferentiated febrile patients were enrolled. Healthy volunteers were recruited to the study from the Faculty of Medicine and Allied Sciences, the Rajarata University of Sri Lanka, to collect control samples upon written consent. Healthy volunteers selected for this study did not exhibit fever-like symptoms, and their white blood cell count (4 - 11 * 10^3/μL) and differential cell count were within the normal range at the time of sample collection.

### Sample collection and storage

A total of 2mL fresh whole blood sample was collected into an EDTA tube by a qualified nursing officer using sterile and standard techniques. 250μL volumes of whole blood were aliquoted into Eppendorf tubes immediately after the sample collection, followed by adding TRIzol LS reagent(Thermo Fisher Scientific, USA) according to the instructions. Blood samples were sent to the Rajarata University of Sri Lanka Public Health Research laboratory to store at − 80^0^C.

### Disease confirmation

Prior to TLR2 studies, disease confirmation was done using previously published qPCR[20] protocol. All qPCR experiments in this study were performed using the CFX96 real-time PCR detection system (Bio-Rad, USA). Total DNA was extracted from patient whole blood samples and healthy whole blood samples, as per the manufacturer’s instructions of the DNeasy kit using the protocol for ‘Purification of Total DNA from Animal blood or cells (Spin-Column Protocol)’ (Qiagen, Germany). DNA qPCR was performed using the primer pair targeting16 S rRNA gene (Forward-5’GCGTAGGCGGACATGTAAGT-3’ Reverse-5’AATCCCGTTCACTACCCACG-3’) [20] with the following thermal cycle conditions:95ᵒ C for 5min,45 cycles of[94ᵒ C for 30s,60 ᵒC for 30s], followed by a melt curve from 65ᵒ C to 90ᵒ C performed at an increment of 0.5 ᵒC per cycle. DNA qPCR was performed in a 20μL reaction mixture containing 10μL of PerfeCTa SYBR Green FastMix(Quanta bio, USA), 0.02μL each of 100μM forward and reverse primers, 4.96μL of nuclease-free water, and 5μL of DNA template, on the CFX96 real-time PCR detection system (Bio-rad, USA).

### Human TLR2 primer designing, validation and selection for gene expression analysis

Two primer pairs of human TLR2 were designed using the NCBI gene database and Primer-BLAST tool based on the available gene sequence of human Toll-like receptor-2 (S1 Appendix and S2 Appendix). Primer pair-I is an exon-exon span primer, which spans an exon junction (exon junction is in between 212/213 bp on the reverse primer) of the mRNA template leads to limiting the amplification only to mRNA, not residual genomic DNA, while primer pair-II was designed to target mRNA transcripts and genomic DNA of the target gene [21,22]. The role and the option availability of designing exon junction span primers were described in the Primer-BLAST primer designing tool on https://www.ncbi.nlm.nih.gov/tools/primer-blast. Primers were verified by *in-silico* PCR on the UCSC web server (https://genome.ucsc.edu/cgi-bin/hgPcr). Gradient PCR was performed to optimize the PCR thermal conditions in the reaction. Agarose gel electrophoresis was performed to confirm the correct amplicon size of the primers (S1 Fig).

### Human reference gene primer designing and validation

Primer pairs (exon junction span primers) of human reference genes were designed using NCBI gene database and Primer-BLAST tool based on the available gene sequences (https://www.ncbi.nlm.nih.gov/tools/primer-blast) (S3 Appendix and S4 Appendix). These primers were designed appropriately as in the instructions of the primer designing for human TLR2 gene. Beta-2-microglobulin exon junction is between 97/98 base pairs of the forward primer and Hypoxanthine phosphoribosyltransferase 1 exon junction is between 174/175 base pairs of the forward primer (Table 2). Primers were verified by *in-silico* PCR on the UCSC web server (https://genome.ucsc.edu/cgi-bin/hgPcr). Gradient PCR was performed to optimize the PCR thermal conditions in the reaction. Agarose gel electrophoresis was performed to confirm the correct amplicon size of the primers(S2 Fig).

**Table 1:**
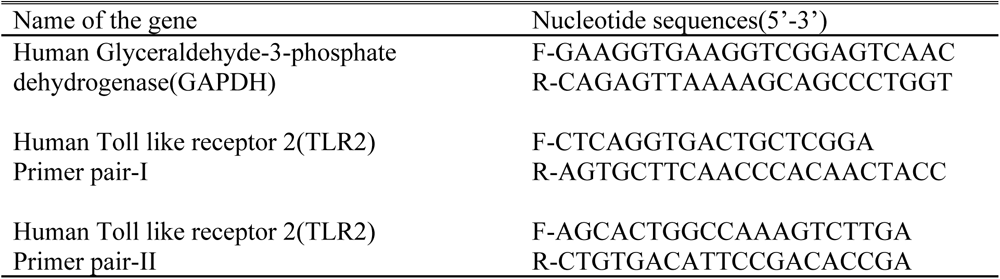
Primer pairs used for Experiment 1.

**Table 2:**
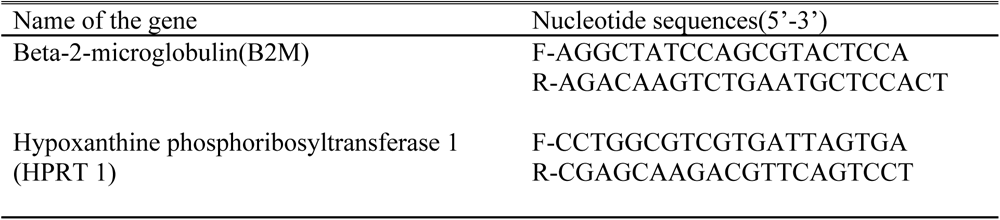
Primer pairs used for Experiment 2.

### Total RNA extraction and quantification

Total RNA was extracted from the RNAlater added samples corresponding to the DNA qPCR positive samples for *Leptospira*, using the manufacture’s guidelines of TRIzol LS reagent(Thermo Fisher Scientific, USA) protocol. The quantity of the RNA was assessed prior to the experiments according to the guidelines of the Qubit RNA high sensitivity (HS) assay kit and Qubit Fluorometer(Thermo Fisher Scientific, USA).

### Complementary DNA (cDNA) synthesis

cDNA was synthesized according to the manufacture’s guidelines of SuperScript™ IV VILO™ Master Mix with ezDNase enzyme kit(Thermo Fisher Scientific, USA). For one gDNA(genomic DNA) digestion reaction, 8μL of total RNA(∼160ng of RNA) was mixed with a gDNA digestion mixture. After the gDNA digestion, RT(reverse transcriptase) and NRT(No reverse transcriptase) reactions were carried out for cDNA synthesis. All incubation steps were performed using the Analytik-Jena thermal cycler (Biometra GmbH, USA). The synthesized cDNA samples were stored at −80^0^C.

### Selection of the most suitable primer pair for the gene expression analysis

Selected sample series was used to perform RT qPCR, by amplifying the target gene using TLR2 primer pair I and II separately. GAPDH primer was used to amplify the reference gene. Non template controls (NTC) were run in duplicates. No Reverse transcriptase (NRT) was run in a single well for each sample. By comparing the expression results of two primer pairs, the most suitable primer pair was selected for the analysis.

### Human TLR2 gene expression analysis

This study was conducted in two separate Experiments with the availability of patient samples.

#### Experiment 1

Leptospirosis confirmed samples and healthy control samples were used to perform the human TLR2 gene expression analysis. GAPDH (Glyceraldehyde-3-phosphate dehydrogenase) was selected as the reference gene/housekeeping gene. GAPDH primer pairs[23] were used. Target gene (human TLR2-primer pair-1), Non Template Controls (NTC) were run in duplicates. No Reverse Transcriptase (NRT) was run in a single well for each sample.

#### Experiment 2

Leptospirosis confirmed samples and healthy control samples were used to perform the human TLR2 gene expression analysis. Human B2M and HPRT 1 were used as the reference gene/housekeeping genes. Target gene (human TLR2-primer pair-1), Non Template Controls (NTC) were run in duplicates. No Reverse Transcriptase (NRT) was run in a single well for each sample.

### Data analysis

Real-time PCR system(qPCR system-CFX Maestro™ software) was selected to compare the human TLR2 gene expression levels among patients confirmed as leptospirosis cases. Further, normalization against a reference gene(housekeeping gene) method was selected to calculate the Relative normalized expression values according to Pfaffl, 2004. Statistical analysis was done by CFX96 Maestro™ software (version 1.1). P-value is calculated using an unpaired t-test.

## Results

### DNA qPCR for Leptospirosis

A total of 100 RNA-stabilized samples were collected from febrile patients with a suspected diagnosis of leptospirosis during the period from October 2018 to December 2018. Among these samples, 64 were subsequently confirmed to be cases of leptospirosis using qPCR. Among the 64 confirmed cases, 18 samples with a leptospirosis quantification cycle (Cq) of less than 40 and a total RNA quantity of 20 ng/μL or more were chosen for gene expression analysis.

For the experiment 2, a total of 80 RNA-stabilized samples were obtained from febrile patients with a probable leptospirosis diagnosis between December 2020 and May 2022. Among these samples, 23 were confirmed as cases of leptospirosis through qPCR. From the 23 confirmed cases, 13 samples meeting the criteria of a Cq value of less than 40 for leptospirosis and a total RNA quantity of 20 ng/μL or more were selected for gene expression analysis.

### Primer optimization and validation

Optimized thermal conditions for human TLR2 gene primers and human reference gene primers are shown in the Human TLR2 PCR products were visualized on a 2% Agarose gel containing ethidium bromide to confirm the correct amplicon size of the primers (S1Fig and S2Fig).

Table 3. Human TLR2 PCR products were visualized on a 2% Agarose gel containing ethidium bromide to confirm the correct amplicon size of the primers (S1Fig and S2Fig).

**Table 3:** Optimized thermal conditions of TLR2 primers and human reference gene primers.

| Primer pair | Thermal Condition |
| --- | --- |
| TLR2<br>Primer pair I | 95°C-2 min of initial denaturation followed by 44 cycles of [95°C for 10 sec, 58°C for 30 sec, 60°C for 30 sec] followed by a melt curve from 65°C to 90°C performed at an increment of 0.50°C per cycle. |
| TLR2<br>Primer pair II | 95°C-2 min of initial denaturation followed by 44 cycles of [95°C for 10 sec, 56°C for 30 sec, 58°C for 30 sec], followed by a melt curve from 65°C to 90°C performed at an increment of 0.5 °C per cycle |
| B2M | 95°C-2 min of initial denaturation followed by 44 cycles of [95°C for 10 sec, 55.1°C for 30 sec, 72°C for 30 sec], followed by a melt curve from 65°C to 90°C performed at an increment of 0.5 °C per cycle |
| HPRT1 | 95°C-2 min of initial denaturation followed by 44 cycles of [95°C for 10 sec, 57.5°C for 30 sec, 72°C for 30 sec], followed by a melt curve from 65°C to 90°C performed at an increment of 0.5 °C per cycle |

### Selection of the most suitable primer pair for the gene expression analysis

Divergent outcomes were observed in the Relative Normalized Expression of the human TLR2 gene when employing Primer Pair I and Primer Pair II on the same set of samples, as evident in Table 4. The disparities in gene expression are visually represented in S3 Fig. Despite the consistent application of DNase treatment to all total RNA samples prior to RT qPCR, it was noted that Primer Pair II exhibited TLR2 gene amplification in six specific samples (A824, A839, A853, A884, A910, and PO47) as well as in the No Reverse Transcriptase (NRT) control reactions for samples A824, A839, A853, A884, and A910. These observations suggest that the utilization of Primer Pair II may introduce errors in gene expression results due to the potential presence of residual genomic DNA. Consequently, for the gene expression analysis, only Primer Pair I was utilized, as it exclusively amplified the mRNA sequence, reducing the risk of genomic DNA interference.

**Table 4:** Comparison of the Relative normalized expression of TLR2.

| Sample | Expression of<br>TLR2(Primer pair I) | Expression of<br>TLR2(Primer pair II) | Regulation(Primer<br>pair-I) | Regulation(Primer<br>pair-II) |
| --- | --- | --- | --- | --- |
| A812 | 0.00000 | 0.00000 | No change | No change |
| A820 | 0.00000 | 0.00000 | No change | No change |
| A822 | 0.00000 | 0.00000 | No change | No change |
| A824 | 0.00000 | 0.17891 | No change | Down regulated |
| A832 | 0.00000 | 0.00000 | No change | No change |
| A833 | 0.00000 | 0.70899 | No change | No change |
| A839 | 0.00000 | 53.61714 | No change | Up regulated |
| A842 | 0.00000 | 0.00000 | No change | No change |
| A853 | 0.00000 | 0.01415 | No change | Down regulated |
| A884 | 0.00000 | 0.00017 | No change | Down regulated |
| A910 | 0.00000 | 0.00390 | No change | Down regulated |
| PO47 | 0.01352 | 0.67494 | Downregulated | Down regulated |

Results of the human TLR2 gene expression are presented separately in two experiments.

#### Experiment 1

Among the 18 samples analyzed, the GAPDH gene was found to be expressed in 13 of them. However, in 5 samples, neither the GAPDH gene nor the TLR2 gene exhibited any expression. Both genes (TLR2 and GAPDH) were expressed by the sample PO47 (Table 5). The normalized gene expression value (ΔΔCq) of the TLR2 of PO47 was 0.01352, showing a ‘downregulation’ of the gene while all others showed a ‘no change’ in gene regulation.

**Table 5:** mRNA expression of TLR2 gene(target),GAPDH gene(Reference), Relative normalized expression of the human TLR2 gene and the Regulation of TLR2.

| Sample | Mean Cq<br>(quantification<br>cycle) of Human<br>TLR2 | Mean Cq<br>(quantification<br>cycle) of Human<br>GAPDH | Relative<br>normalized<br>expression($\Delta\Delta Cq$)<br>of TLR2 | Regulation of<br>TLR2 |
| --- | --- | --- | --- | --- |
| A809 | 0 | 22.75 | 0.00000 | No change |
| A812 | 0 | 00.00 | 0.00000 | No change |
| A820 | 0 | 00.00 | 0.00000 | No change |
| A822 | 0 | 00.00 | 0.00000 | No change |
| A824 | 0 | 30.44 | 0.00000 | No change |
| A829 | 0 | 24.72 | 0.00000 | No change |
| A832 | 0 | 00.00 | 0.00000 | No change |
| A833 | 0 | 24.24 | 0.00000 | No change |
| A834 | 0 | 30.75 | 0.00000 | No change |
| A839 | 0 | 32.79 | 0.00000 | No change |
| A842 | 0 | 00.00 | 0.00000 | No change |
| A853 | 0 | 24.21 | 0.00000 | No change |
| A884 | 0 | 27.81 | 0.00000 | No change |
| A888 | 0 | 22.53 | 0.00000 | No change |
| A903 | 0 | 37.38 | 0.00000 | No change |
| A910 | 0 | 24.71 | 0.00000 | No change |
| PO47 | 37.98 | 29.25 | 0.01352 | Down regulated |
| PO77 | 0 | 26.95 | 0.00000 | No change |
| CONTROL | 27.42 | 24.90 | Not applicable | Not applicable |

#### Experiment 2

Both genes (target and reference) were expressed in 3 out of 13 samples (Table 6 and Table 7). Normalized gene expression value (ΔΔCq) of the TLR2 (reference B2M) were 0.86574, 0.47200 and 0.28579 in S1, S6 and S8 respectively. ΔΔCq of the TLR2 (reference HPRT1) were 0.48226, 0.11121, 0.12349 in S1, S6 and S8 respectively. Across all samples, a ‘downregulation’ was noted when applying a two fold change threshold for both reference genes. However, when the threshold was increased to fourfold, the B2M gene expression remained unchanged. For HPRT1, samples S6 and S8 exhibited ‘downregulation’, whereas sample S1 displayed ‘no change’ at the fourfold change threshold.

**Table 6:** mRNA expression of TLR2 gene(target), B2M gene(Reference), Relative normalized expression of the human TLR2 gene and the Regulation of TLR2.

| Sample | Mean Cq<br>(quantification<br>cycle) of<br>Human TLR2 | Mean Cq<br>(quantification cycle)<br>of Human HPRT1 | Relative<br>normalized<br>expression($\Delta\Delta Cq$)<br>of TLR2 | Regulation of<br>TLR2 |
| --- | --- | --- | --- | --- |
| S1 | 32 | 33.00 | 0.48226 | Downregulation |
| S2 | 0 | 00.00 | 0.00000 | No change |
| S3 | 0 | 00.00 | 0.00000 | No change |
| S4 | 0 | 00.00 | 0.00000 | No change |
| S5 | 0 | 00.00 | 0.00000 | No change |
| S6 | 38 | 37.00 | 0.11121 | Downregulation |
| S7 | 0 | 00.00 | 0.00000 | No change |
| S8 | 32 | 31.00 | 0.12349 | Downregulation |
| S9 | 0 | 00.00 | 0.00000 | No change |
| S10 | 0 | 00.00 | 0.00000 | No change |
| S11 | 0 | 00.00 | 0.00000 | No change |
| S12 | 0 | 00.00 | 0.00000 | No change |
| S13 | 0 | 00.00 | 0.00000 | No change |
| CONTROL | 29 | 32.00 | 1.00000 | Not applicable |

**Table 7:** mRNA expression of TLR2 gene(target), HPRT1 gene(Reference), Relative normalized expression of the human TLR2 gene and the Regulation of TLR2.

| Sample | Mean Cq<br>(quantification<br>cycle) of<br>Human TLR2 | Mean Cq<br>(quantification cycle)<br>of Human B2M | Relative<br>normalized<br>expression($\Delta\Delta Cq$)<br>of TLR2 | Regulation of<br>TLR2 |
| --- | --- | --- | --- | --- |
| S1 | 32 | 23.00 | 0.86574 | Downregulation |
| S2 | 0 | 00.00 | 0.00000 | No change |
| S3 | 0 | 00.00 | 0.00000 | No change |
| S4 | 0 | 00.00 | 0.00000 | No change |
| S5 | 0 | 00.00 | 0.00000 | No change |
| S6 | 38 | 28.00 | 0.47200 | Downregulation |
| S7 | 0 | 00.00 | 0.00000 | No change |
| S8 | 32 | 21.00 | 0.28579 | Downregulation |
| S9 | 0 | 00.00 | 0.00000 | No change |
| S10 | 0 | 00.00 | 0.00000 | No change |
| S11 | 0 | 00.00 | 0.00000 | No change |
| S12 | 0 | 00.00 | 0.00000 | No change |
| S13 | 0 | 00.00 | 0.00000 | No change |
| CONTROL | 29 | 21.00 | 1.00000 | Not applicable |

### Visualization of the gene expression results

The Relative normalized expression of the human TLR2 gene in experiment 1 (S4 and S5 Fig) and 2 (S6 and S7 Fig) were analyzed using CFX Maestro™.

## Discussion

The study aimed to investigate the human TLR2 gene’s response to leptospiral infection using RT qPCR. Primer Pair 1, which specifically amplifies mRNA sequences of TLR2, was selected to avoid non-specific genomic DNA amplification. Among the 31 samples, 4 samples showed TLR2 downregulation, while the rest did not yield normalized gene expression values.

This study’s primary strength lies in using human samples from confirmed cases of leptospirosis instead of in-vitro or animal studies to study the TLR2 response within a real biological system. Additionally, the use of the RT qPCR assay to determine mRNA levels of the TLR2 gene expression also enhances the validity of our results [25,26]. mRNA transcripts represent the activated genes of the organism during a particular time period. Hence, the assay demonstrates the gene expression results more accurately [24]. Even though we did the DNase treatment prior to RT qPCR assay, residual genomic DNA would be amplified eventually in the reaction plate. Though the use of exon-exon spanning primer is debate [21,27], to get rid of gDNA contaminants, we designed and validated an exon-exon spanning primer pair that specifically amplifies the mRNA sequences only. By using an exon junction span primers we could avoid false positive results.

Despite having at least more than ∼160 ng of total RNA in the cDNA (complementary DNA) reaction well, 27 out of 31 samples in our experiment did not show normalized gene expression value (ΔΔCq). mRNA is fragile and easily degradable [28] and collection of clinical samples requires RNA stabilization at the time of sample collection. Though we followed the RNA stabilization procedure, the lack of gene expression in the majority of our samples shows the degradation of mRNA has occurred in these samples, limiting the sample size with generalizable results. As in most other gene expression studies [29], we were unable to collect samples in the early stage of the disease, which is unavoidable due to the ubiquitous nature of the disease. It is also difficult to collect samples prior to antibiotic treatments as patients usually come to the hospital after initial treatment, and this may alter the innate immune responses and our results.

In contrast to findings of mice models and *in vitro* studies, in our study, we found four samples showed downregulation of the human TLR2 gene during leptospirosis. To our knowledge, there is no human study evidence reported on the downregulation of TLR2 gene during the infection of *Leptospira*. Previous studies using flowcytometry showed increased TLR2 expression on human polymorphonuclear cells[30] and human neutrophils[31], while one study showed no significant TLR2 expression difference [32] on monocytes of confirmed patients for leptospirosis compared to healthy human samples. With the different types of methodologies used, detection of expression of the human TLR2 gene can be varied. Flowcytometry directly focuses the receptors expressed on immune cells, while RTqPCR assay targets mRNA transcripts only. However, the observed differences could not be attributed only to the differences in methodologies used.

Nevertheless, studies on other bacterial infections demonstrated the downregulation of TLR2 gene via different types of mechanisms. Kim and the colleagues [33] demonstrated that TLR2 downregulation may be associated with patient mortality in the early stage of *Staphylococcus aureus* bacteremia (SAB). Several other studies on *Staphylococcus aureus* have also demonstrated the human and mouse TLR2 inactivation during the bacterial infection[34–36]. This process is mediated through Staphylococcal superantigen-like protein (SSL3) that directly binds to the extracellular domain of TLR2 and inhibits TLR2 activation on human and murine monocytes, macrophages and neutrophils. Though the exact role of SSL3 on TLR2 inhibition has not been described; inactivation of TLR2 was detected by the reduced level of cytokine secretion in response to interaction of SSL3 vs TLR2. Current knowledge of the interaction between human TLR2 vs SSL3 has been expanded to reveal the molecular basis interaction of the both components[37]. Whether a similar mechanism exists in *Leptospira* infection is yet to be investigated.

Micro RNAs (miRNAs) are a novel group of small RNA molecules that firstly described in nematodes 1993[38,39]. These molecules do not encode proteins, but regulate the gene expression in humans. About 1% of human genome acquired by miRNAs, regulate up to 60% of all protein-coding genes[40]. miRNAs can inhibit the target gene expression or can degrade the target gene mRNA transcript by base pairing to the 3’ untranslated region(UTR) of the target gene mRNA[41]. Syphilis is caused by the bacterium *Treponema pallidum*, that mounts the immune responses in humans but the regulation of the human immune responses caused by the bacterium is not clearly defined. One study has suggested the infection has up-regulated the human Micro RNA, miR-101-3p and it paired with the 3’ UTR of mRNA of the TLR2 gene to downregulate the TLR2 gene expression and eventually reduced the cytokine secretion[42]. Not like bacterial infections, virus itself can express Micro RNA s during human infection, for instance, miR-UL112-3p has been demonstrated as a human TLR2 downregulator. TLR2 downregulation inhibits the downstream cascade of reactions, eventually leads to evade the antiviral immune responses[43]. Accordingly, pathogen secreting components and micro RNA can down regulate the TLR2 expression during infection.

Recognition of PAMPs by TLR2 mediates the secretion of pro-inflammatory cytokines that defend against bacterial infection. Excessive production of inflammatory cytokines may result in tissue damage, sepsis, and eventually death. Likewise, TLR2 downregulation may help reduce cytokine levels, prior to an excessive cytokine secretion(cytokine storm). Zhaoxia Haung and the group [44] have synthesized a tripalmitoylated lipopeptide (Pam3CSK4) to use as a pretreatment against Methicilin-resistant *Staphylococcus aureus* (MRSA). Pam3CSK4 is synthetic but similar ligand with bacterial lipoproteins that binds to TLR1/TLR2 receptor. Pretreated mice with Pam3CSK4 showed downregulation of TLR2 gene and down-stream molecules IRAK-1(Interleukin-1 receptor Associated kinases) during the bacterial infection. Pam3CSK4 was used in an in-vivo leptospirosis study[45] in order to stimulate TLR2. Knowing the accurate response level of TLR2, is worth to regulate the level of immune effectors secrete during leptospirosis using newly invented therapeutics.

Even though in-vivo and in-vitro studies have suggested the increase of TLR2 expression during leptospirosis, only a few human studies performed the TLR2 response against leptospirosis. It is worth to increase the number of human studies to be performed on TLR2 response against the infection of *Leptospira* spp. When doing the experiments it is important to collect samples at the early stage of disease onset and prior to antibiotic treatments. It is reliable to perform the experiment as soon as possible to make sure that the mRNA fragments are not degraded due to long term storage even after the RNA stabilization. As reported in other bacterial diseases, blocking of TLR2 by bacterial components or TLR2 mRNA degradation by micro RNA need to be clarified. Our study’s key finding is the downregulation of the TLR2 gene expression in human leptospirosis, challenging previous assumptions of TLR2 upregulation in in-vivo and in-vitro studies. This observation diverges from the anticipated upregulation and underscores the need for a revised understanding of TLR2’s role in leptospirosis pathogenesis. The use of exon-exon spanning primers provides a more precise approach for gene expression analysis and should be considered best practice in future research. These insights into TLR2 gene behavior offer a foundation for potential advancements in diagnostic and therapeutic strategies for leptospirosis.

## Acknowledgements

We would like to thank Joseph M. Vinetz and Michael A. Matthias at School of Medicine, Yale University, New Haven, Connecticut, United States of America. We extend our thank to Nursing Officer Ms. K.S.Shashini Kaushalya and the technical staff at Rajarata University’s Public Health Research Laboratory, Faculty of Medicine and Allied Health Sciences, Rajarata University of Sri Lanka. We also express our gratitude to Professor Kosala Weerakoon from the Department of Parasitology, alongside the dedicated physicians and healthcare staff at the Teaching Hospital Anuradhapura, for their indispensable contributions to our work.

## Supporting information

S1 Appendix. Primer Designing steps-Human TLR2 primer pair 2

S2 Appendix. Primer Designing steps-Human TLR2 primer pair 1

S3 Appendix. Primer Designing steps-Human HPRT1

S4 Appendix. Primer Designing steps-Human B2M

S1 Figure. Visualized PCR bands of human TLR2 primer products on a 2% Agarose gel containing Ethidium bromide

S2 Figure. Visualized PCR bands on a 2% Agarose gel containing Ethidium Bromide

S3 Figure. Relative normalized expression of TLR2 amplified using Primer pair I (right hand side) and Primer pair II (left hand side)

S4 Figure. The Relative normalized expression of human TLR2 gene (PO47 showed expression)

S5 Figure. The Relative normalized expression of human TLR2 gene (PO47 showed expression)

S6 Figure. The Relative normalized expression of human TLR2 gene (reference B2M)S1,S6, S8

S7 Figure. The Relative normalized expression of human TLR2 gene (referenceHPRT1) S1,S6,S8

## Availability of data, code and other materials

All the data produced or examined during this study have been incorporated in the published article (along with supplementary materials).

